# Antibiotic resistance is lower in *Staphylococcus aureus* isolated from antibiotic-free raw meat as compared to conventional raw meat

**DOI:** 10.1101/448209

**Authors:** Haskell Kyler J., Schriever Samuel R., Fonoimoana Kenisi D., Haws Benjamin, Hair Bryan B., Wienclaw Trevor M., Holmstead Joseph G., Barboza Andrew B., Berges Erik T., Heaton Matthew J., Bradford K. Berges

**Affiliations:** Department of Microbiology and Molecular Biology, Brigham Young University Provo, UT 84602, USA; Department of Statistics, Brigham Young University, Provo, UT 84602, USA

**Keywords:** *Staphylococcus aureus*, MRSA, Antibiotic resistance, Antibiotic abuse, Food microbiology, Livestock

## Abstract

The frequent use of antibiotics contributes to antibiotic resistance in bacteria, resulting in an increase in infections that are difficult to treat. Livestock are commonly administered antibiotics in their feed, but there is current interest in raising animals that are only administered antibiotics during active infections. *Staphylococcus aureus* (SA) is a common pathogen of both humans and livestock raised for human consumption. SA has achieved high levels of antibiotic resistance, but the origins and locations of resistance selection are poorly understood. We determined the prevalence of SA and MRSA in conventional and antibiotic-free (AF) meat products, and also measured rates of antibiotic resistance in these isolates. We isolated SA from raw conventional turkey, chicken, beef, and pork samples and also from AF chicken and turkey samples. We found that SA contamination was common, with an overall prevalence of 22.64% (range of 2.78-30.77%) in conventional meats and 13.0% (range of 12.5-13.2%) in AF poultry meats. MRSA was isolated from 15.72% of conventional raw meats (range of 2.78-20.41%) but not from AF-free meats. The degree of antibiotic resistance in conventional poultry products was significantly higher vs AF poultry products for a number of different antibiotics, and while multi-drug resistant strains were relatively common in conventional meats none were detected in AF meats. The use of antibiotics in livestock contributes to high levels of antibiotic resistance in SA found in meat products. Our results support the use of AF conditions for livestock in order to prevent antibiotic resistance development in SA.

## Introduction

The discovery of antibiotics has saved countless lives as they have been used to treat bacterial infections. However, bacteria can quickly develop resistance to antibiotics through mutation and by horizontal gene transfer [1]. Many bacterial species have acquired resistance to a number of antibiotics and the rate of development of new antibiotics is not keeping pace with the development of resistance. High rates of antibiotic use by humans, by livestock animals, and also the release of antibiotics into the environment continue to select for resistant hosts [2]. As a result, many bacterial infections are difficult to treat and future prospects are not promising that the trend will reverse.

Livestock animals are commonly raised in high density environments; thus infectious agents rapidly move through animals resulting in significant morbidity/mortality. These animals are commonly administered antibiotics prophylactically to prevent bacterial infections. Prophylactic use of antibiotics results in better animal survival, and also in higher meat yields. This practice is widespread around the world, and current estimates suggest that 80% of all antibiotics are administered to livestock [3]. This high rate of antibiotic use can result in the development of antibiotic resistance in livestock-associated bacterial species. Since many bacteria that infect livestock also infect humans (e.g., *E. coli, S. aureus, Salmonella*), the areas where livestock are raised are thought to be a breeding ground for antibiotic resistance [4].

*Staphylococcus aureus* (SA) is an opportunistic bacterial pathogen carried asymptomatically by healthy individuals; it is found consistently in 20% and intermittently in 60% of the human population [5]. SA can carry a number of virulence genes, including hemolysins, enterotoxins, and immune-modulatory factors [6, 7]. SA can cause a variety of human diseases including skin infections, sepsis, and pneumonia [7-9]. It is also likely the most common cause of food poisoning in the United States [10].

Methicillin-resistant *Staphylococcus aureus* (MRSA) is a group of SA strains that has become resistant to many common antibiotics (methicillin-susceptible strains referred to as MSSA). SA and MRSA have become an increasing problem in healthcare in the United States, where they cause an estimated 80-100,000 invasive infections and 11-19,000 deaths per [11, 12]. In the U.S., the Centers for Disease Control and Prevention concluded that during the year 2012, the number of MRSA-infected patients admitted into the hospital was estimated to be just above 75,000 [13]. The most common mode of SA/MRSA transmission is person-to person contact, and transmission usually takes place either in hospitals or the community [14-20]. However, some individuals become infected with SA/MRSA through live animal contact, and others through contact with raw livestock meat [16-20]. These bacteria can experience horizontal gene transfer by mobile genetic elements that confer antibiotic resistance, which is thought to be the cause of the emergence of resistant bacteria found in farms and farm workers [21]. There is a link between voluntary removal of antibiotics from large-scale farms and a significant reduction in rates of antibiotic resistance among *Enterococcus* isolates from those farms [22]. These data suggest that antibiotic resistance in livestock can be reversible.

The aim of this study was to determine the prevalence of SA and MRSA in raw beef, chicken, pork, and turkey meats from conventionally-raised animals, and also from chicken and turkey meats from antibiotic-free (AF) raised animals, and to characterize individual antibiotic resistance profiles of the isolates. Data obtained were then analyzed to determine correlations of meat types and levels of antibiotic resistance to see if there were differences in rates of antibiotic resistance in SA isolated from meats of a particular species. We also determined if there were differences in rates of antibiotic resistance in meats from animals raised with or without prophylactic antibiotic use.

## Materials and Methods

### Isolation and Identification of SA/MRSA in meat samples

Conventional meat samples were collected from at least 11 different grocery stores/wholesale stores/ethnic markets (see Table 2), which were obtained as packaged meats at grocery and wholesale stores and unpackaged meats from ethnic markets. AF meat samples were collected from 5 different stores, representing 6 different brands (see Table 3), all as packaged meats. Samples were tested for SA by swabbing meat with a sterile swab or pipetting 10pl of meat juice directly onto Mannitol Salt Agar (MSA) plates. MSA plates that showed no growth were scored as negative for SA. Growth on MSA plates, accompanied by fermentation, initially indicated SA detection. Gram stains were performed to confirm the presence of gram-positive cocci, and all isolates were also catalase and coagulase positive. Genotyping was performed by PCR to detect the presence of Staphylococcus-specific 16S rDNA sequences, and *nucA* detection was used to confirm *S. aureus* [23]. MRSA was detected by the same procedure in the presence of 2 pg/mL oxacillin. To confirm MRSA detection, PCR was used to detect *mecA* [23]; products were separated on a 1.5% agarose gel.

### Disk Diffusion Test

The disk diffusion test was used to classify the resistance of each isolate to antibiotics. We used a standard protocol [24] and used ATCC *S. aureus* reference strain 25923 as a control; if that strain failed to show established resistance values then the test was repeated. Mueller-Hinton agar plates were used for growth, supplemented with 2% NaCl. McFarland standards were used to verify that bacterial density was in the appropriate range. Amounts of antibiotic per disk were as follows: clindamycin 2 μg, cefotaxime 30 μg, gentamicin 10 μg, erythromycin 15 gg, tetracycline 30 gg, ciprofloxacin 5 μg, chloramphenicol 30 μg, and rifampin 5 μg (Sigma Aldrich). Susceptibility of isolates were identified by zone diameters as determined by CLSI standards. Plates were incubated at 37°C for 24-48 hours.

### Minimum Inhibitory Concentration (MIC)

MIC tests were performed for vancomycin at concentrations of 6 μg/mL (intermediate resistance) and 16 μg/mL (complete resistance). Mueller-Hinton agar plates were prepared with vancomycin, then inoculated with isolates and analyzed for growth after 24-48 hours at 37°C. MRSA was detected by the same procedure in the presence of 4 gg/mL oxacillin. Plates were observed to see if growth occurred; if only a few colonies were detected, then samples were retested. Presence of just a few colonies was not scored as resistant.

### Statistical analysis

To analyze the prevalence of SA, MSSA, or MRSA in different types of raw meat samples, a chi-squared test was performed. A two sample t-test with unequal variance was used to determine if the sum of all meat isolates was more or less resistant/susceptible to a certain antibiotic. For all antibiotics tested with zone diameter measurements, t-tests were performed, while a 2-sample z-test of proportions was performed on oxacillin and vancomycin and also for differences in the prevalence of MDR strains. Multi-drug comparison was performed by an asymptotic chi-square test. A Fishers Exact test was used to examine differences in rates of antibiotic resistance to oxacillin and vancomycin.

## Results

### Detection and isolation of SA in conventional raw meat samples

159 different conventional meat samples (beef, chicken, pork and turkey) were tested for SA, as described in Materials and Methods. 1 of 36 beef samples was positive (2.78%), 14 of 49 chicken samples were positive (28.57%), 12 of 39 pork samples were positive (30.77%), and 9 of 35 turkey samples were positive (25.71%). SA contamination in beef was significantly lower than all other meat types (p<0.001; chi-squared test), but there were no other significant differences in prevalence. The overall frequency of SA isolation from the 159 samples was 36 (22.64%; see Table 1). The meat samples were collected from at least 11 different stores, as summarized in Table 2. Information on the store of origin was not available for all 36 isolates, but the origin is reported for 20 of the 36 isolates (55.6%).

**Table 1.** Prevalence of *Staphylococcus aureus* in Raw Meat Samples

| Meat type | Number of samples tested | Number of SA isolated | % with SA | Number of MSSA isolated | % with MSSA | Number of MRSA isolated | % with MRSA |
| --- | --- | --- | --- | --- | --- | --- | --- |
| C Beef | 36 | 1 | 2.78% | 0 | 0.00% | 1 | 2.78% |
| C Chicken | 49 | 14 | 28.57% | 4 | 8.16% | 10 | 20.41% |
| C Pork | 39 | 12 | 30.77% | 5 | 12.82% | 7 | 17.95% |
| C Turkey | 35 | 9 | 25.71% | 2 | 5.71% | 7 | 20.00% |
| C Total | 159 | 36 | 22.64% | 11 | 6.92% | 25 | 15.72% |
| AF Chicken | 53 | 7 | 13.21% | 7 | 13.21% | 0 | 0% |
| AF Turkey | 24 | 3 | 12.50% | 3 | 12.50% | 0 | 0% |
| AF Total | 77 | 10 | 13.00% | 10 | 13.00% | 0 | 0% |

Prevalence of *Staphylococcus aureus* (SA) in raw meat samples, which is further divided into Methicillin-Susceptible *Staphylococcus aureus* (MSSA) and Methicillin-Resistant *Staphylococcus aureus* (MRSA). C=Conventional meat sample (raised with antibiotics) and AF=Antibiotic-free meat sample.

**Table 2.** Origin of *Staphylococcus aureus* Isolates (Conventional Meats) by Store Location

| Store | Beef<br>MRSA | Chicken<br>MSSA | Chicken<br>MRSA | Pork<br>MSSA | Pork<br>MRSA | Turkey<br>MSSA | Turkey<br>MRSA | Total |
| --- | --- | --- | --- | --- | --- | --- | --- | --- |
| Grocery store A |  |  | 1 |  |  |  |  | 1 |
| Grocery store B |  | 1 |  |  |  |  |  | 1 |
| Grocery store C |  | 1 | 1 |  |  |  |  | 2 |
| Grocery store D |  |  | 1 |  |  |  |  | 1 |
| Grocery store E |  |  |  | 2 | 1 | 2 | 1 | 6 |
| Grocery Store F |  |  |  |  | 1 |  |  | 1 |
| Wholesale store A |  |  | 1 |  |  |  | 1 | 2 |
| Wholesale store B |  |  | 2 | 1 |  |  |  | 3 |
| Small ethnic market<br>A |  |  | 1 |  |  |  |  | 1 |
| Small ethnic market<br>B |  |  |  |  | 1 |  |  | 1 |
| Small ethnic market<br>C |  |  |  | 1 |  |  |  | 1 |
| Unknown origin* | 1 | 2 | 3 | 1 | 4 |  | 5 | 16 |
| Total | 1 | 4 | 10 | 5 | 7 | 2 | 7 | 36 |

Store locations and specific isolates per store are listed to show that SA isolates were collected from diverse locations. *Some meat samples were provided without any information on the store of origin.

### Screening of SA isolates from conventional meats for MRSA

SA isolates were re-plated on MSA plates in the presence of 2 pg/ml oxacillin to determine resistance to oxacillin. Any isolates found to be resistant to oxacillin were initially classified as MRSA, and all others were determined to be MSSA. To confirm MRSA, PCR genotyping was performed (data not shown) to detect the *mecA* gene; all isolates reported as MRSA produced a band of ~533bp (see Methods). MRSA was detected only rarely in beef (1 of 36 meat samples positive; 2.78%), but was common in the other three meat types: 10 of 49 meat samples positive in chicken (20.41%), 7 of 39 meat samples positive in pork (17.95%), and 7 of 35 meat samples positive in turkey (20.00%). The overall frequency of MRSA isolation from the 159 samples was 15.72% (Table 1), consistent with reports by others in different locations [25]. Of samples where SA was detected, 100% were MRSA in beef (but n=1), 77.78% were MRSA in turkey, 71.43% were MRSA in chicken, and 58.33% were MRSA in pork. Overall, the majority of SA isolates were MRSA (69.44%). Beef had significantly fewer MSSA (p<0.05) and MRSA (p<0.02) contamination as compared to other meat types, but there were no other significant differences by meat type.

### Antibiotic resistance in conventional raw meat SA isolates

We next measured antibiotic resistance in all MSSA and MRSA isolates. Resistance levels were determined by disk diffusion for eight common antibiotics. Supplemental Table 1 shows disk diffusion distances for each isolate, with zone diameters (in millimeters) for all antibiotics except for oxacillin and vancomycin. Relative antibiotic resistance (complete resistance, intermediate resistance, and complete susceptibility) is also indicated for each isolate. We detected two isolates with intermediate resistance to vancomycin, one of which was also a MRSA strain. Mean disk diffusion distances are shown in Figure 1 to better illustrate relative differences by meat type and by antibiotic. Of note, the frequency of isolates that were multidrug resistant (MDR; complete resistance to three or more antibiotics) per meat group were found to be: 100% (beef; but n=1), 66.7% (chicken), 55.5% (turkey), and 54.5% (pork); the overall frequency of multi-drug resistance was 20/33 or 60.6%. No significant differences in multi-drug resistance were found by meat type; beef was excluded due to the small sample size. Complete susceptibility to all antibiotics tested was not detected in any isolate.

**Figure 1:**
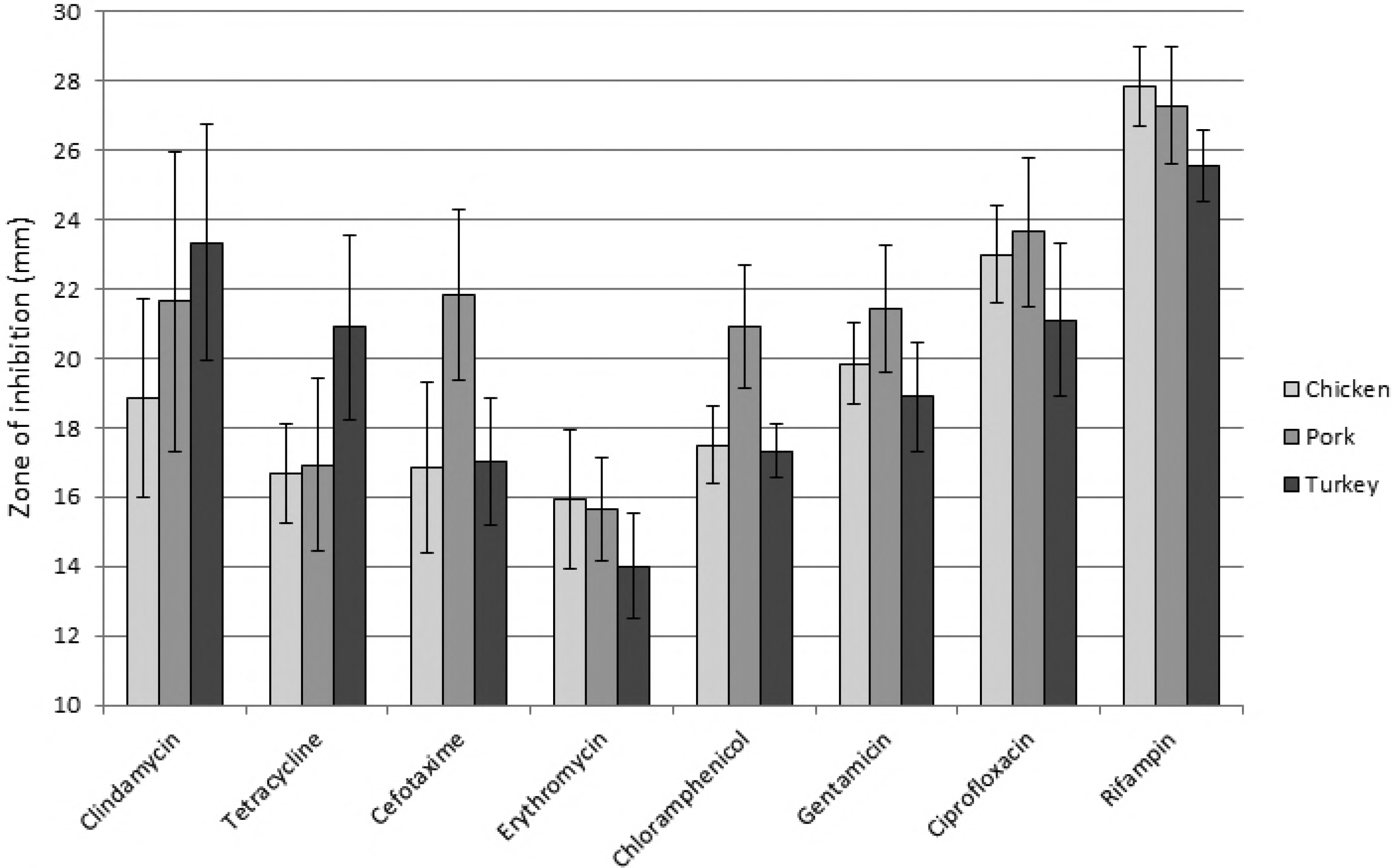
Mean antibiotic resistance in conventional raw meat SA isolates for eight common antibiotics. Disk diffusion tests were performed to determine the amount of antibiotic required to prevent growth of the various SA raw meat isolates. There was only a single SA isolate from beef, and thus no results are reported here for that meat type. Means were calculated and standard error is indicated. Antibiotic concentrations used and significance of zone diameters are detailed in Methods. A two sample t-test with unequal variance test was used to determine if there were significant differences in rates of antibiotic resistance or susceptibility amongst the various meat types.

We then determined if there were any significant differences in rates of antibiotic resistance when comparing meat types. The only significant result was that pork SA isolates were significantly more susceptible to cefotaxime as compared to other SA isolates (p=0.03 for susceptibility, or p=0.97 for resistance). We compared rates of antibiotic resistance amongst all SA samples to determine if SA from raw meat samples showed significant differences in antibiotic susceptibility across the 10 antibiotics tested (Fig. 1). Clindamycin resistance was significantly higher that rifampin (p=0.005); tetracycline resistance was significantly higher than rifampin (p<0.001) and ciprofloxacin (p=0.003); cefotaxime resistance was significantly higher than rifampin (p<0.001) and ciprofloxacin (p=0.011); erythromycin resistance was significantly higher than all other antibiotics (p<0.05); chloramphenicol resistance was significantly higher than rifampin (p<0.001) and ciprofloxacin (p=0.002); gentamicin resistance was significantly higher than rifampin (p<0.001) and ciprofloxacin (p=0.035); and ciprofloxacin resistance was significantly higher than rifampin (p=0.001).

### Detection and isolation of SA and MRSA in AF raw poultry samples

Raw meat samples were tested for the presence of SA using the same methods outlined above, but using raw meat samples marked as “antibiotic-free”. In total, 77 different raw poultry meat samples (chicken and turkey) were tested. AF meat sources are shown in Table 3. 7 of 53 chicken samples were positive for SA (13.2%) and 3 of 24 turkey samples were positive (12.5%) for SA. SA was found at significantly higher levels in conventional meats as compared to AF meats for chicken (p=0.03), but not for turkey (p=0.11). Isolates were genotyped as above to confirm SA and MRSA. None of the 10 isolates from AF poultry meats were positive for the *mecA* gene, indicating a lack of MRSA amongst all AF poultry isolates. These results were further confirmed by a lack of growth on MSA plates with 2 pg/ml oxacillin. MRSA was found at significantly higher levels in conventional meats as compared to AF meats for both chicken (p=0.0004) and turkey (p=0.0002).

**Table 3.** Origin of *Staphylococcus aureus* Isolates (Antibiotic-free meats) by Store Location

| Store | Chicken<br>MSSA | Chicken<br>MRSA | Turkey<br>MSSA | Turkey<br>MRSA | Total |
| --- | --- | --- | --- | --- | --- |
| Grocery store A, Brand 1 | 2 | 0 | 2 | 0 | 4 |
| Grocery store B, Brand 1 | 5 | 0 | 1 | 0 | 6 |
| Grocery store C, Brand 1 | 0 | 0 | 0 | 0 | 0 |
| Grocery store D, Brand 1 | 0 | 0 | 0 | 0 | 0 |
| Grocery store E, Brand 1 | 0 | 0 | 0 | 0 | 0 |
| Grocery store E, Brand 2 | 0 | 0 | 0 | 0 | 0 |
| Total | 7 | 0 | 3 | 0 | 0 |

Store locations and specific isolates per store are listed to show that SA isolates were collected from diverse locations.

### Antibiotic resistance in AF poultry SA isolates

We next measured antibiotic resistance in all SA isolates from AF meats, as above for conventional meat SA isolates. Supplemental Table 2 shows disk diffusion distances for each isolate, with zone diameters (in millimeters) for all antibiotics except for oxacillin and vancomycin; relative rates of antibiotic resistance are also indicated for each isolate. Mean disk diffusion distances are shown in Figure 2. We did not detect any MDR isolates in the AF meat samples. Complete susceptibility to all antibiotics tested was detected in one chicken SA isolate. All AF SA isolates were susceptible to chloramphenicol, oxacillin, and vancomycin.

**Figure 2:**
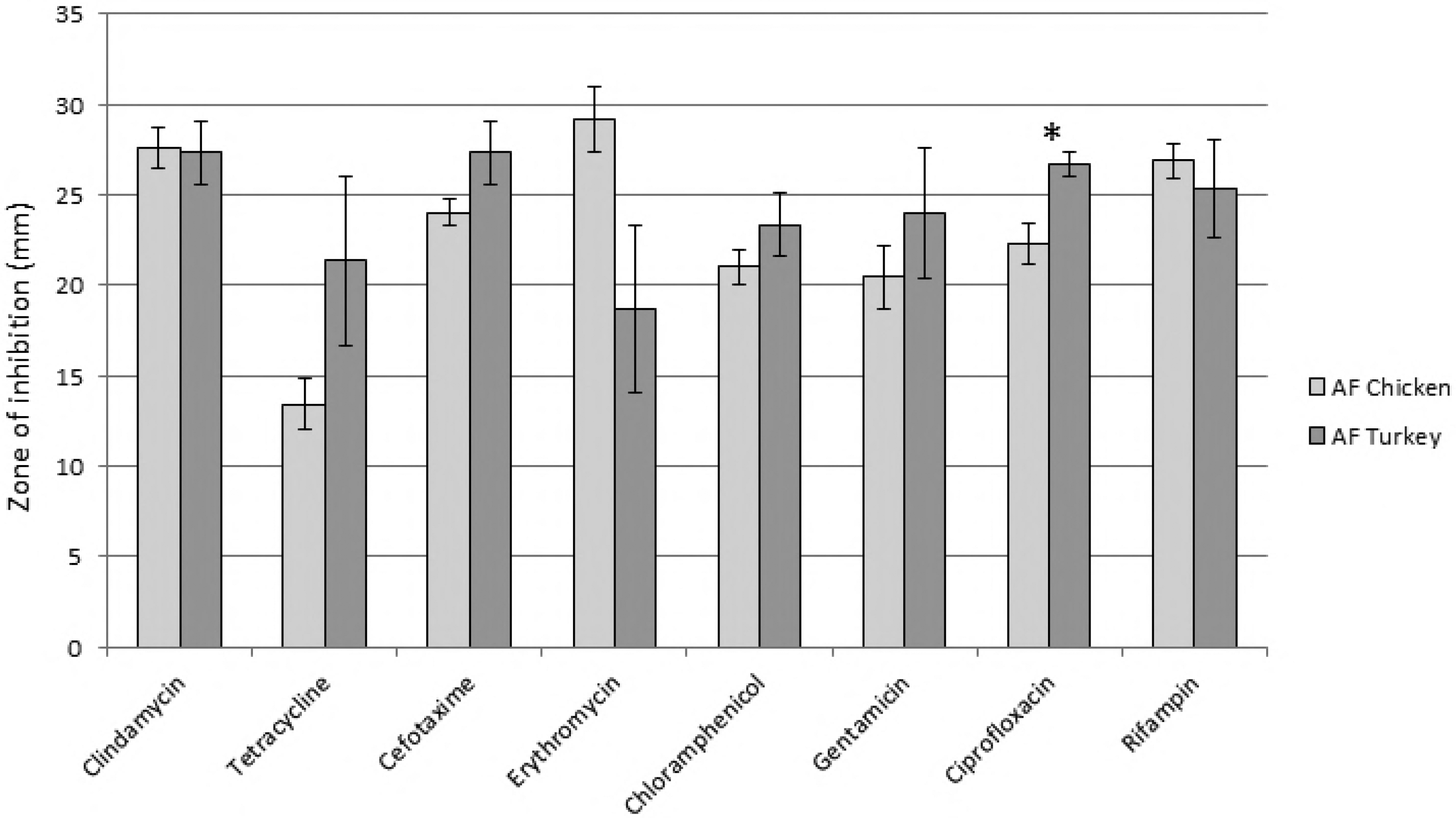
Mean antibiotic resistance in AF raw poultry SA isolates for eight common antibiotics. Disk diffusion tests were performed to determine the amount of antibiotic required to prevent growth of the various SA raw meat isolates. A two sample t-test with unequal variance test was used to determine if there were significant differences in rates of antibiotic resistance or susceptibility amongst the various meat types. * indicates p<0.05.

Statistical analysis was then performed to determine if there were any significant differences in antibiotic resistance when comparing AF poultry meat types. Most antibiotic resistances were not significantly different when comparing SA from chicken or turkey, with the exception being that SA from chicken was significantly more resistant to ciprofloxacin (p=0.005). Chicken isolates approached significantly higher resistance to cefotaxime (p=0.07), and turkey isolates approached significantly higher resistance to erythromycin (p=0.07); a larger sample size for AF turkey isolates might yield significant results for those groups.

Statistical analysis was also performed to determine if there were any significant differences in antibiotic resistance when comparing conventional to AF meat sources (Figure 3). We found significantly lower rates of resistance (p<0.05) to the following antibiotics in AF chicken products: clindamycin, cefotaxime, erythromycin, chloramphenicol, and oxacillin (Fig. 3A). In addition, there was a highly significant difference (p=0.0003) in MDR strains, with none detected in AF chicken meat products. We found significantly lower rates of resistance (p<0.05) to the following antibiotics in AF turkey products: cefotaxime, chloramphenicol, ciprofloxacin, and oxacillin (Fig. 3B). There was a significant difference (p=0.0067) in MDR strains, with none detected in AF turkey meat products.

**Figure 3:**
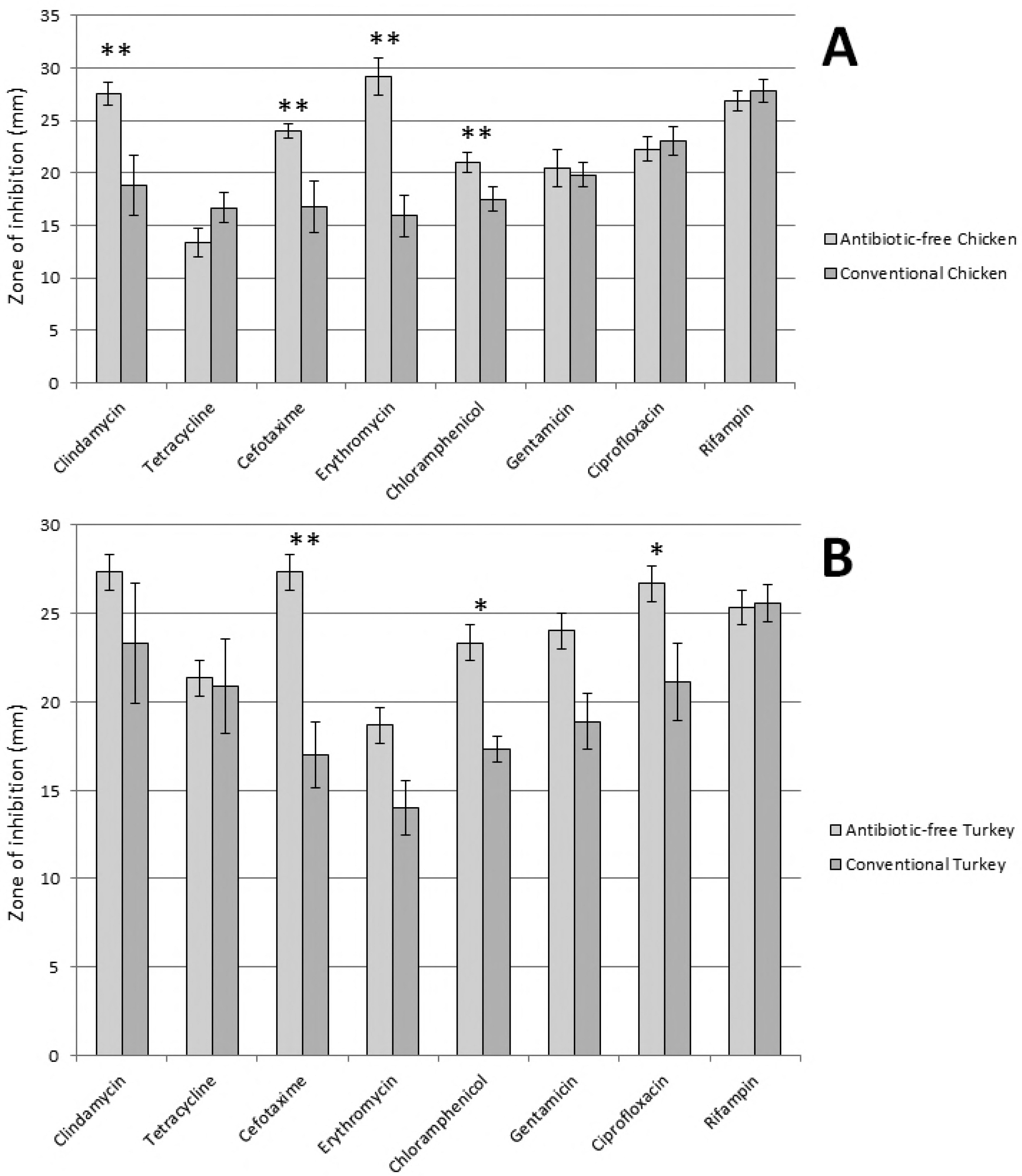
Antibiotic resistance levels in AF meat products as compared to conventional meat products. Panel A shows differences in antibiotic resistance in SA isolated from AF chicken products (n=7) as compared to conventional chicken products (n=12) and panel B shows differences in rates of antibiotic resistance in SA isolated from AF turkey products (n=3) as compared to conventional turkey products (n=9). A 2-sample z-test on proportions was used to determine if there were significant differences in rates of antibiotic resistance amongst the various meat types for all antibiotics tested by disk diffusion. A Fishers Exact test was used to examine significant differences in oxacillin and vancomycin resistance. * indicates p<0.05; ** indicates p<0.01.

## Discussion

We isolated 36 *Staphylococcus aureus* (SA) strains from conventional raw meat products and 10 SA strains from AF meat products. SA was common in conventional raw meat products, with a combined prevalence of 22.6% amongst the four meat types. Beef contamination was significantly lower than other meat types, but no significant differences were seen between non-beef frequencies. MRSA isolates were also common in conventional meat products, with an overall prevalence of 15.7%, but there were no significant differences in either MRSA or MSSA detection with the exception that beef had significantly lower contamination with both types. SA contamination of AF poultry meats was significantly lower than in conventional meats (13.0% vs 22.6%; p=0.02), and no MRSA was detected in the 77 AF poultry samples (0%; p<0.001). We also determined the antibiotic resistance profiles of each isolate for ten common antibiotics. Antibiotic resistance in SA was very common amongst conventional meat isolates, but less common in AF meat isolates. 20 conventional meat isolates showed resistance to at least three different antibiotics (60.6% of the isolates); while no isolates were multi-drug resistant in the AF group.

The prevalence of SA detected in our conventional meat samples was remarkably consistent amongst chicken, pork and turkey (range of 25.71-30.77%). We detected more MRSA than MSSA in raw meat samples (Table 1), and that trend held true across all meat types. Our MRSA detection was higher than reported in other areas of the USA, especially for MRSA in poultry, but were lower than those reported for pork in Canada [26]. Our AF meats had a significantly lower SA prevalence than for conventional meats. The reasons for this finding are not clear, but since SA has high rates of antibiotic resistance it is possible that this species can outcompete other species when antibiotics are present, but when they are not it is outcompeted by other bacteria due to higher fitness in other areas.

It is possible that contamination of meat products at central processing locations could explain our results. Our isolates were obtained from at least 11 different stores, and the antibiotic resistance profiles shown provide evidence that we did not re-isolate the same strains repeatedly because the drug resistance patterns amongst the various isolates match only rarely (Suppl. Tables 1 and 2). Taken together, this analysis indicates that many independent SA strains were isolated during these studies, suggesting that a common source of SA contamination at a processing plant is less likely to have affected our results. This further supports the hypothesis that SA and MRSA isolates obtained from raw consumer meats include a variety of SA strains with different resistance profiles, which could contribute to eventual increased resistance in strains that could become a concern to consumers due to potential mobile genetic elements.

The United States Food and Drug Administration releases results of the total amounts of antibiotics sold for use in food-producing animals, and the 2014 report showed an increase of 22% in sales from 2009 to 2014. Tetracycline accounted for 70% of sales, followed by penicillin (9%), macrolides (7%), and sulfas (5%) with no other drugs over 3% [27]. We have shown that tetracycline resistance is significantly higher in SA isolates from conventional meats than that seen for rifampin or ciprofloxacin, but not for other the antibiotics tested (Figure 1), although there was no significant difference. High levels of antibiotic usage in livestock can also be seen on a worldwide scale; in 2014 it was estimated that 38.5 million kilograms of antibiotics were used exclusively in swine and poultry in China [28] and it is estimated that worldwide antibiotic usage will increase 67% from 2010 to 2030 due to growing demand for meat [29].

A handful of studies have analyzed the frequency of SA and MRSA in meats produced from animals raised under AF conditions. A study of AF vs conventional chicken products in Oklahoma, USA found a lower prevalence of SA contamination in AF meats vs. conventional meats (41% vs 53.8%) but the difference was not significant. MRSA contamination was very low in both types of chicken [30]. SA was somewhat less common in AF pork (56.8%) as compared to conventional pork (67.3%), although the difference was not significant. MRSA frequency in raw pork was very similar in conventional pork (6.3%) and AF pork (7.4%) [31]. *E. coli* antibiotic resistance in organic vs conventionally-raised pigs in four European countries was found to be lower for a number of different antibiotics tested [32], and analysis of antimicrobial resistance genes across the microbiome of AF vs conventional chickens found that AF animals also had lower levels of antibiotic resistance genes in their associated bacteria [33]. These results indicate that the lower rates of antibiotic resistance in AF animals likely apply to other bacterial species and not just for SA. A poultry farm in the USA transitioned from common antibiotic use to organic practices, and two *Enterococcus* species were analyzed for antibiotic resistance before and after the transition. Interestingly, antibiotic resistance levels significantly decreased following the transition [22].

## Conclusions

In conclusion, we have found that the use of antibiotics in livestock contributes to high levels of antibiotic resistance in SA isolates found in their resulting meat products. Our results suggest that the use of antibiotics in livestock promotes higher rates of antibiotic resistance in bacteria found in the meat products that consumers come into contact with and could be a source of transmission of antibiotic resistance bacteria to humans. AF conditions for livestock may prevent antibiotic resistance development in SA and in other microbes, and could relieve the continued development of antibiotic resistance.

## List of Abbreviations

SA: Staphylococcus aureus
MSSA: Methicillin-susceptible *Staphylococcus aureus*
MRSA: Methicillin-resistant *Staphylococcus aureus*
LA: livestock-associated.

## Acknowledgements

We thank Kyle Jensen, Taalin Rasmussen, Kelsey Berges, and Chloe McCullough for help in collecting meat samples.

